# Analysis of factor V in zebrafish demonstrates minimal levels needed for hemostasis and risk stratifies human variants

**DOI:** 10.1101/567453

**Authors:** Angela C. Weyand, Steve. J. Grzegorski, Megan. S. Rost, Kari. I. Lavik, Allison C. Ferguson, Marzia Menegatti, Catherine E. Richter, Rosanna Asselta, Stefano Duga, Flora Peyvandi, Jordan A. Shavit

## Abstract

In humans, coagulation factor V (F5) deficiency is a rare, clinically heterogeneous bleeding disorder, suggesting that genetic modifiers may contribute to disease expressivity. Complete loss of mouse F5 results in early lethality. Zebrafish possess many distinct advantages including high fecundity, optical clarity, external development, and homology with the mammalian hemostatic system, features that make it ideal for genetic studies. Our aim was to study the role of F5 in zebrafish through targeted mutagenesis, and apply the model to the study of human *F5* variants. CRISPR-mediated genome editing of the zebrafish *f5* locus was performed, generating mutants homozygous for a 49 base pair deletion in exon 4. Thrombus formation secondary to vascular endothelial injury was absent in *f5*^*-/-*^ mutant embryos and larvae. Despite this severe hemostatic defect, homozygous mutants survived before succumbing to severe hemorrhage in adulthood. Human *F5* variants of uncertain significance from patients with F5 deficiency were evaluated, and the causative mutations identified and stratified by their ability to restore thrombus formation in larvae. Analysis of these novel mutations demonstrates variable residual F5 function, with minimal activity being required to restore hemostasis. This *in vivo* evaluation may be beneficial for patients whose factor activity levels lack correlation with bleeding symptomatology. Furthermore, homozygous mutant embryos tolerate what is a severe and lethal defect in mammals, suggesting the possibility of species-specific factors enabling survival, and allowing further study not possible in the mouse. Identification of these factors or other genetic modifiers could lead to novel therapeutic modalities.

**Key Points:**

- F5 mutant fish embryos tolerate symptoms lethal in mammals but succumb to bleeding in adulthood
- Analysis of human variants demonstrate that all have some residual function and that minimal F5 activity is required to restore hemostasis

## INTRODUCTION

Coagulation factor V (F5) is an essential component of normal hemostasis. In plasma, it circulates as a large 330 kilodalton glycoprotein that shares homology and a similar domain organization with factor VIII^1,2^. The inactive pro-cofactor form of F5 is synthesized in the liver as a 2240 amino acid polypeptide and after proteolytic removal of the large B domain, it is secreted as a 2196 amino acid single chain protein, circulating in the plasma at a concentration of 20 nmol/L^3–5^. In its activated form, F5a serves as the cofactor protein for the activated serine protease factor Xa (F10a) in the prothrombinase complex, which rapidly converts prothrombin to thrombin. The gene for F5 is located on the long arm of human chromosome 1q23, consists of 25 exons and is approximately 80 kilobases (kb) in length^6^.

Congenital deficiency of F5 in humans, also known as parahemophilia, is a rare autosomal recessive bleeding disorder that affects 1 in 1,000,000 individuals^7–10^. Over 200 cases have been described with over 100 DNA alterations reported, including insertions, deletions, splice site, missense, and nonsense mutations. Two thirds of all mutations causing F5 deficiency have been found to be nonsense mutations in the *F5* gene. Of the missense mutations which have been reported, none have been found in the B domain. Most of these mutations affect secretion of F5 resulting in substantially reduced antigen levels. The spectrum of mutations is quite diverse, with most patients carrying private mutations. F5 deficiency is divided into Type I (quantitative) and Type II (qualitative) defects, with the majority of mutations falling in the former group and approximately 25% in the latter^11^. Current treatment options are limited as only recently has a purified F5 concentration been developed^12^. Additionally, multifactor concentrates such as cryoprecipitate, prothrombin complex concentrates (PCCs) and activated PCCs contain inadequate concentrations of F5 to be effective. For major bleeding, fresh frozen plasma or platelet transfusions are typically used^7^, although volume overload is a concern. For minor bleeding, anti-fibrinolytics are often sufficient^9^.

Insight into the crucial role of F5 has been further facilitated using gene targeting in mice. *F5* knockouts exhibit complete lethality, with death occurring in either the embryonic or perinatal periods^13^, although transgenic expression of very small amounts (<0.1%) of *F5* has been shown to rescue this phenotype^14^. In humans, unlike hemophilia A and B, levels of F5 antigen have not been found to correlate well with bleeding symptomatology. Clinically, a wide spectrum exists. In patients with severe F5 deficiency, typically with F5 levels below 10 to 15%, mucosal bleeding is seen most commonly, affecting approximately 60% of patients. Fewer patients present with menorrhagia, hematomas and hemarthroses^15^. The relatively mild bleeding seen in many patients with severe F5 deficiency may be explained by findings that <1% F5 is adequate to ensure minimal thrombin generation as demonstrated in *in vivo*^16^, *in vitro*^17^, and *in silico*^18^ studies. However, this does not explain the striking variability in phenotype between patients with equivalent or undetectable F5 levels.

Zebrafish (*Danio rerio*) have been used extensively to study hemostasis and have demonstrated significant homology with the mammalian coagulation system. They possess many distinct advantages including high fecundity, optical clarity and external development. New developments in genome editing technology have facilitated quick and robust systems for targeted genetic modification^19–21^. We and others have demonstrated the benefits of this system for modeling hemostasis and thrombosis^22–34^. Knockout models have demonstrated longer than expected survival in comparison to their mammalian counterparts^25,35,36^, enabling additional characterization of the individual phenotypes. We previously demonstrated a severe hemostatic defect in F10 deficient zebrafish very early in development with lack of venous occlusion in response to endothelial injury at 3 days post fertilization (dpf)^36^. Homozygous mutants were able to tolerate this defect with no overt hemorrhage until 3-4 weeks of age, and survival of the majority for several months. Anticoagulant deficiency has also been shown to exhibit less severe phenotypes in the zebrafish in comparison to murine models. A zebrafish model of antithrombin (At3) deficiency exhibited disordered coagulation with a resulting phenotype similar to clinical disseminated intravascular coagulation^25^. Spontaneous thrombi and consumption of fibrinogen led to increased clotting times in larvae, while adult mutants ultimately succumbed to extensive intracardiac thrombosis. Despite rampant dysregulation of coagulation, homozygous mutants lived to adulthood, in stark contrast to the prenatal mortality observed in a murine model^25,37^. Here we report targeted ablation of zebrafish *f5* using the CRISPR/Cas9 genome editing platform. Homozygous mutant embryos and larvae demonstrate a severe defect in hemostasis. They tolerate this into adulthood, but the majority eventually succumb to lethal bleeding by 6 months of age. The ability to adapt to what is a severe and early lethal defect in mammals suggests the possibility of unique fish species specific hemostatic factors and also allows further studies not possible in the mouse knockout.

## METHODS

### Zebrafish strains and maintenance

Zebrafish were raised in accordance with animal care guidelines as approved by the University of Michigan Animal Care and Use Committee. Embryos are defined as 0 to 2 dpf, larvae 3 to 29 dpf, juvenile 30 to 89 dpf, and adult ≥90 dpf^38^. *f5* mutant zebrafish were generated on an AB X TL hybrid background.

### Targeted mutagenesis of the *f5* locus using CRISPR/Cas9 genome editing

The *f5* locus was identified in the zebrafish genomic sequence assembly^39^ and CRISPR/Cas9 mediated genome editing was used to induce mutations in exon 4. Target sites of 17-20 nucleotides were identified using the web based ZiFiT Targeter program (http://zifit.partners.org) as described^40^. Templates for synthetic single guide RNAs (sgRNAs) were constructed using PCR to fuse overlapping oligonucleotides (Integrated DNA Technologies, Coralville, IA) which contained the complementary 20 nucleotides of the target site, the T7 promoter, and linearized vector pDR274^41^ as a template (Table 1). The resulting double stranded DNA fragment (120 base pairs, bps) was then column purified and transcribed using the T7 Quick High Yield RNA Synthesis kit (New England BioLabs, Ipswich, MA) and purified using RNA Clean and Concentrator kit (Zymo Research, Irvine, CA). Cas9 mRNA was transcribed as previously described^21^. sgRNAs and Cas9 mRNA were injected concurrently into the cytoplasm of one cell stage embryos at concentrations of 12.5 and 300 ng/μL, respectively. The resulting F0 population was raised to adulthood and crossed with wild type zebrafish to verify germline transmission to the F1 generation.

**Table 1:**
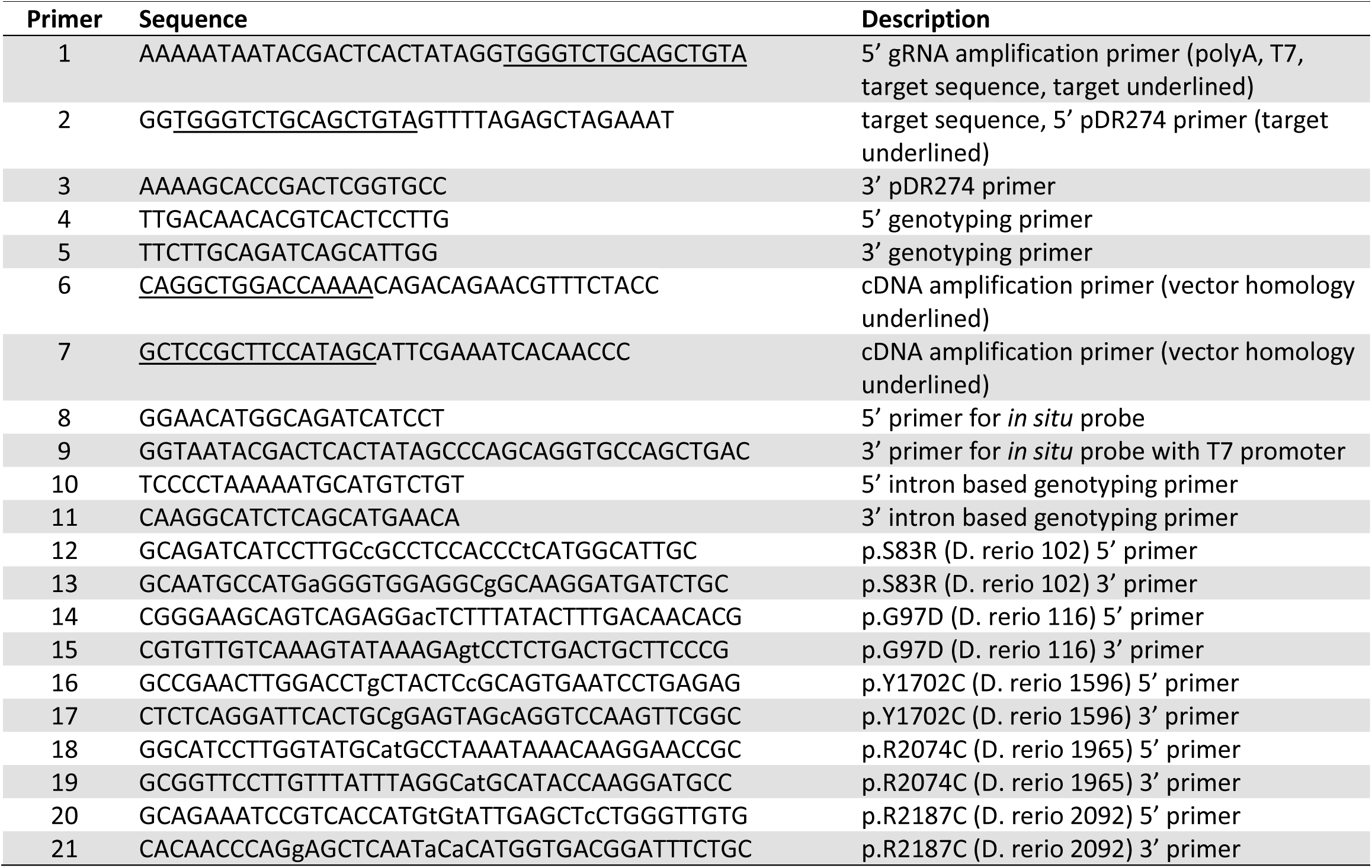
Primers used in experimental procedures

### Genotyping of *f5* mutant offspring

Staged zebrafish larvae or adults were anesthetized in tricaine (0.16 mg/mL, Western Chemical Inc., Ferndale, WA) or humanely killed in high dose tricaine (1.6 mg/mL). Fish were tail fin-clipped as described^25,42^. Genomic DNA was isolated by incubating tissue in lysis buffer (10 mM Tris-Cl, pH 8.0; 2 mM EDTA, 2% Triton X-100, and 100 μg/mL proteinase K) at 55°C overnight, followed by proteinase K inactivation at 95°C for five minutes^25^. Insertion and deletion mutations were detected by PCR and resolved by agarose gel electrophoresis or using a Qiaxcel (Qiagen, Hilden, Germany). Primers were designed using Primer3^43^ and oligonucleotides ordered from Integrated DNA Technologies (Table 1).

### Laser-induced endothelial injury in zebrafish larvae

Laser-mediated endothelial injury was performed on the posterior cardinal vein (PCV) of zebrafish larvae at 3 dpf as described^26^. Larvae were anesthetized in tricaine, embedded in 0.8% low melt agarose on glass cover slips, and the endothelium of the PCV at the 5^th^ somite distal to the anal pore was ablated using 100 pulses at power level 18 (MicroPoint Pulsed Laser System, Andor Technology, Belfast, Northern Ireland). Arterial injury was also performed on the dorsal aorta at the 5^th^ somite distal to the anal pore at 9 dpf and endothelial injury to the primordial midbrain channel (PMBC) just lateral to the otic capsule at 6-7 dpf. The time to occlusion was recorded for up to 2 minutes by a blinded observer. Larvae were subsequently recovered from agarose and genotyped.

### Construction of *f5 in vivo* expression vectors

pDestTol2pA2_*ubi*:p2A-EGFP^36^ contains a BbsI restriction site flanked multiple cloning site and a self-cleaving p2A peptide 5’ to the *egfp* sequence. The vector was linearized using BbsI and purified. The *D. rerio f5* complementary DNA (cDNA) was amplified from plasmid MDR1734-202728340 (Dharmacon, Lafayette, CO) using primers with *f5* and vector homology (Table 1). The resulting 6.3 kb product was purified and inserted into BbsI linearized pDestTol2pA2_ubi:p2A-EGFP using In-Fusion Cloning (Clontech, Mountainview, CA) and transformed into high efficiency 10-beta competent *E.coli* (New England BioLabs) to generate pDestTol2pA2_*ubi:f5-p2A-EGFP*. Positive colonies were grown up in LB media supplemented with 50 μg/mL ampicillin and sequence verified. Mutations were introduced using the QuickChange XL II site-directed mutagenesis kit (Agilent) and mutation specific primers (Table 1). Amino acid numbering reflects the human amino acid position.

### One-cell injection of *f5* expression vectors

Circular *f5* expression plasmids (10 ng/μL) and transposase mRNA (50 ng/μL)^44^ were co-injected into 1-cell stage embryos from *f5*^Δ49+/ -^in-crosses. Fluorescent embryos were selected at 3 dpf for laser-mediated endothelial injury.

### Whole mount in situ hybridization (WISH)

Inhibition of pigmentation in embryos and larvae was accomplished using 0.2 mM phenylthiourea (PTU, Sigma-Aldrich, St. Louis, MO) at 24 hours post fertilization. Staged larvae were fixed in 4% paraformaldehyde (PFA) in phosphate buffered saline with 0.01% Tween-20 (PBST) at 4°C overnight. They were then dehydrated stepwise into 100% ethanol. Partial *f5* cDNA fragments were amplified to synthesize sense and antisense digoxin-labeled RNA riboprobes (DIG RNA labeling kit, Roche, Basel, Switzerland) (Table 1). Amplification of cDNA templates, synthesis of riboprobes and WISH were performed as previously described.^25,45,46^

### Chemical treatment of embryos

Embryos were incubated in 100 mM ε-aminocaproic acid (Sigma-Aldrich) dissolved in system water starting at 24 hours post fertilization and continuing until 3 dpf. At 3 dpf, they were evaluated by laser-mediated endothelial injury.

## RESULTS

### Targeted disruption of *f5* using genome editing nucleases results in spontaneous hemorrhage and adult lethality

To create a model of F5 deficiency, we used the CRISPR-Cas9 genome editing system to target *f5*. Several sgRNAs were designed and constructed to target exon 4 of the zebrafish *f5* genomic locus. The most robust sgRNA resulted in somatic mutations at a frequency of 66.4% (93 of 140 sequenced clones from 2 pools of 8 F0 injected embryos). The resulting embryos were raised to adulthood and 40 F0 founders were mated to wild type fish to verify germline transmission. We identified many potential founders and verified one for further analysis, a 49 bp deletion (*f5*^Δ49^ hereafter referred to as *f5*^-^) within the predicted A1 domain (Figure 1A, 1B). The mutation was confirmed both at the genomic and complementary DNA (cDNA) levels (Figure 1C, D). Heterozygotes from each line were incrossed, and genotyping prior to 1 month of age revealed a normal Mendelian distribution. Beginning at 28 dpf we observed a statistically significant loss of homozygotes, and by 7 months post fertilization (mpf) approximately 90% died (p<0.0001) (Figure 2A, B). Interestingly we saw a significant loss of heterozygotes starting around 18 months of age (p=0.006). Spontaneous hemorrhage was observed grossly in homozygous mutants beginning around 28 dpf, coincident with the beginning of observed lethality (Figure 2C, 2D).

**Figure 1.**
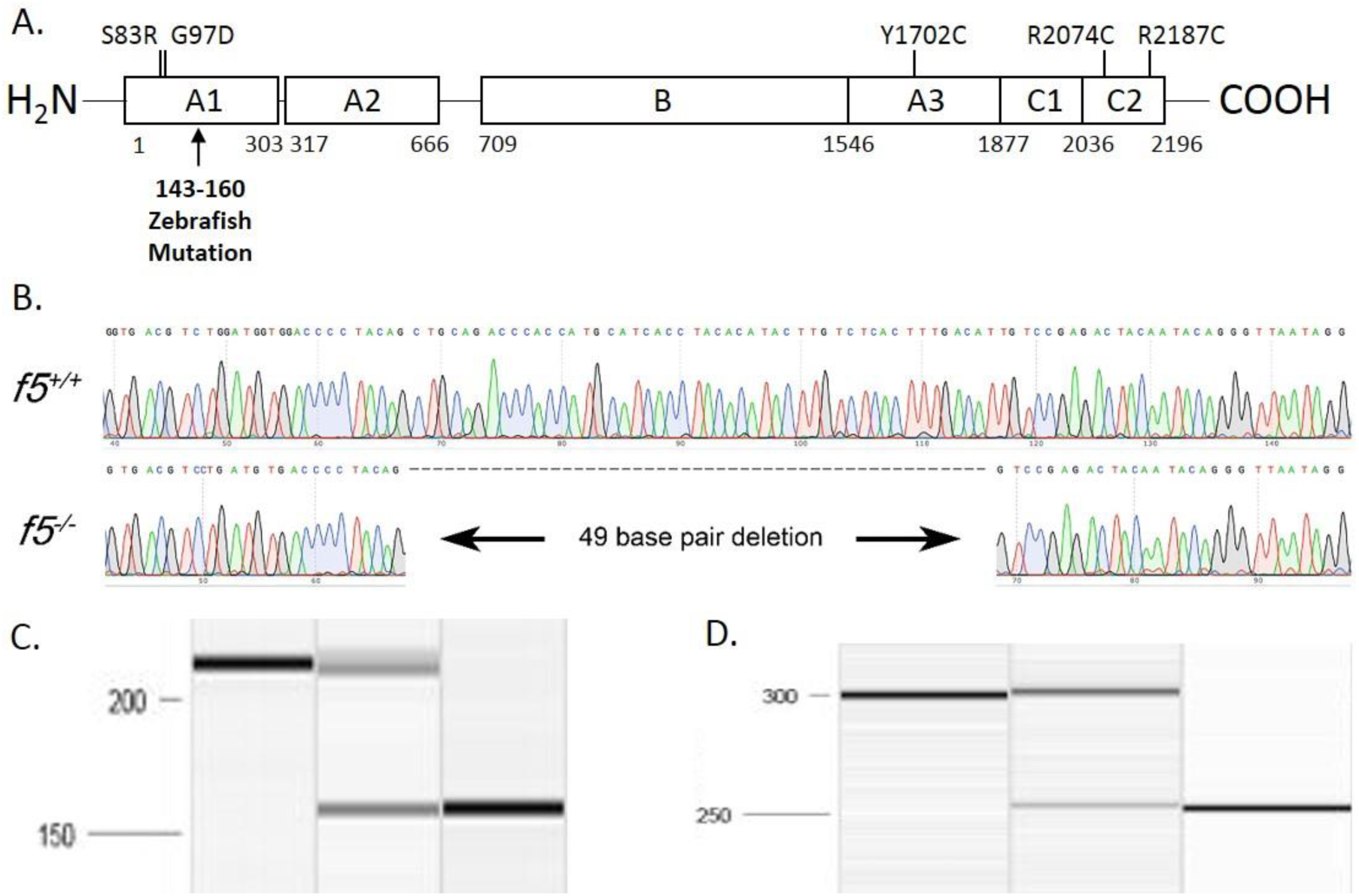
Targeting of the *f5* locus using genome editing produces a null allele. A. Structure of human F5, residue numbering is from the human protein. The CRISPR/Cas9 induced mutation deletes what would be human residues 143-160 in the A1 domain and induces a frameshift. Numbered positions indicate location of human mutations evaluated in Figure 5. B. Sequence of 49 bp *f5* exon 4 deletion. C, D. Amplification of genomic and cDNA from *f5*^*+/+*^, *f5*^*+/-*^, and *f5*^*-/-*^ fish demonstrates the 49 bp deletion with no evidence for any other products.

**Figure 2.**
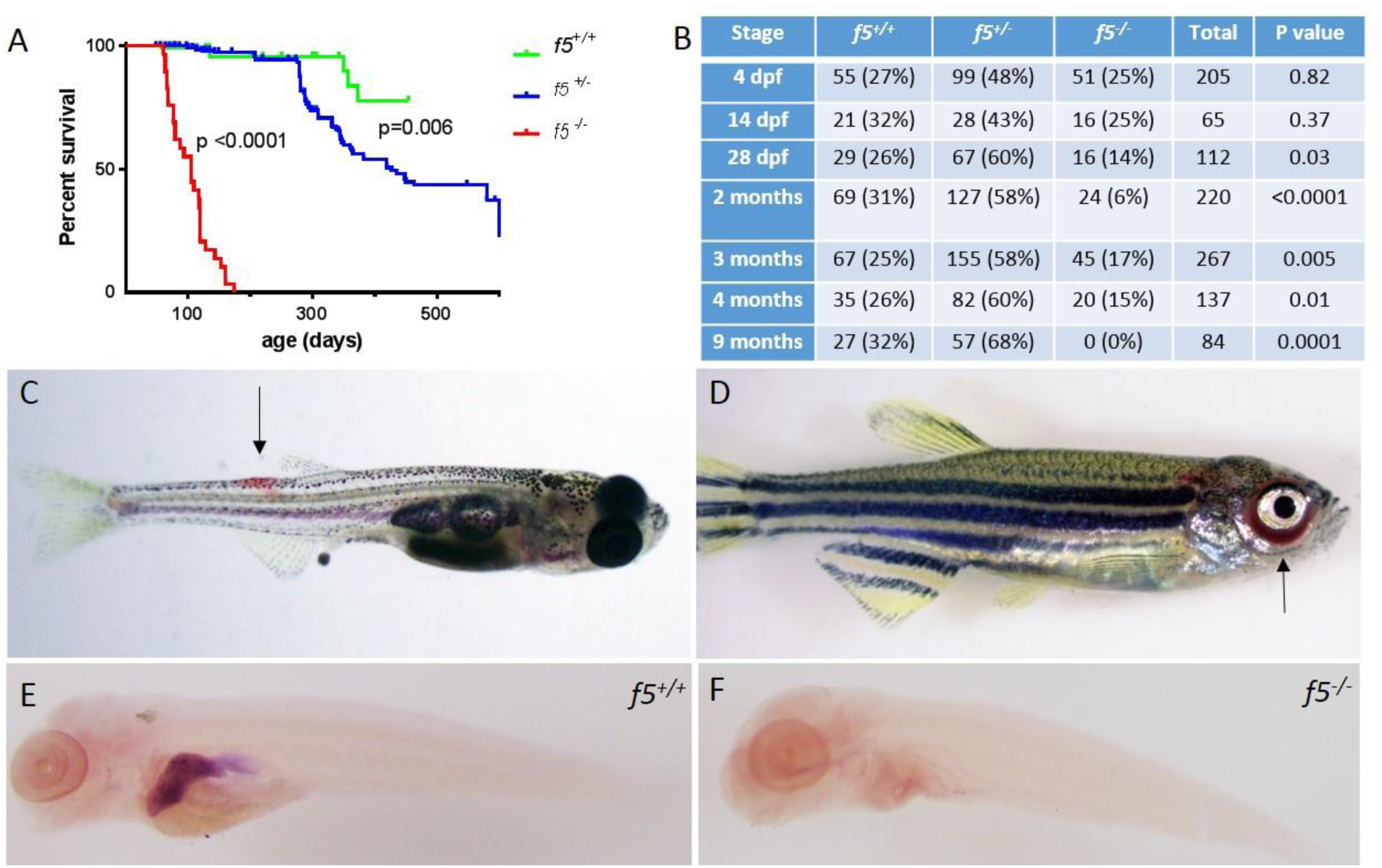
Phenotypes of *f5* homozygous mutants reveal adult lethality and spontaneous hemorrhage in homozygous mutant adult fish. A. Survival curves of zebrafish offspring from *f5*^*+/-*^ incrosses, starting at 2 months of age demonstrate progressive loss of 90% of homozygotes by 7 mpf. Heterozygotes also demonstrate statistically significant decreased survival compared to wild type fish, with 50% mortality by 1.5 years of age. B. Genotype distributions of offspring from *f5*^*+/-*^ incrosses evaluated at various stages demonstrate loss of homozygotes begining at 28 dpf. C. Spontaneous hemorrhage distal to the dorsal fin (arrow) in a 4 week old homozygous mutant fish. D. Severe spontaneous periorbital hemorrhage (arrow) grossly visible in a 5 week old homozygous mutant fish just prior to death. Similar bleeding was not observed by blinded observers in heterozygous or wild type clutchmates. E. Whole mount *in situ* hybridization with an antisense probe demonstrates localization of *f5* expression to the liver in *f5*^*+/+*^ zebrafish at 5 dpf, but not in *f5*^*-/-*^ mutants (F).

### Zebrafish embryos and larvae show no signs of overt hemorrhage

Loss of F5 in mice results in bimodal mortality with half of homozygous mutant embryos dying at embryonic day 9-10, hypothesized to be due to abnormalities in yolk sac vasculature that result in bleeding, and the remaining from massive hemorrhage within two hours of birth^13^. Surprisingly, no appreciable loss of homozygous mutant fish embryos was observed until 28 dpf (Figure 2A). Given embryonic hemorrhage observed in the murine model, we carefully examined zebrafish at early stages of development. Larvae from heterozygous incrosses were stained with o-dianisidine at 7 dpf and examined for any evidence of hemorrhage. No differences in o-dianisidine staining was seen between homozygous mutants and their wild type and heterozygous clutchmates (data not shown). Given the unexpected extended survival and lack of hemorrhage in early embryos and larvae, we hypothesized that there might be residual expression or altered localization of *f5* mRNA in mutants. In zebrafish, the liver bud is first distinguishable at approximately 2 dpf, overlying the yolk sac to the left of midline^47^. The liver grows to its final dimensions between 2.5-3 dpf with outgrowth complete by 5 dpf. Expression of *f5* was examined in the developing zebrafish at 5 dpf, and we confirmed localization to the liver, but it was absent in *f5*^*-/-*^ mutants (Figure 2E, F).

### *f5* homozygous mutants are unable to form thrombi in an induced model of venous and arterial endothelial injury

Since there was no spontaneous bleeding visible during the embryonic/larval period, we evaluated whether *f5* mutants respond to induced trauma. We performed laser-mediated endothelial injury on larvae generated from *f5*^*+/-*^ incrosses, targeting the PCV (orthologous to the mammalian inferior vena cava) at 3 dpf, the dorsal aorta (orthologous to the mammalian aorta) at 9 dpf, and the primordial midbrain channel (PMBC) at 6-7 dpf. As expected, all wild type and *f5*^+/-^ larvae exhibited complete vessel occlusion within two minutes, whereas *f5*^*-/-*^ mutants failed to occlude (venous p=<0.0001, arterial p=0.002, PMBC p=0.008 by the Mann Whitney *U* test, Figure 3A-C). To confirm the role of F5 deficiency in this phenotype, as well as to rule out any potential off target mutations generated through genome editing, we injected *f5*^+/-^ incrosses with a zebrafish *f5* cDNA under control of the zebrafish ubiquitin (*ubi*) promoter (pDestTol2pA2_*ubi:f5-p2A-EGFP*).^48^ Subsequent injury of the PCV at 3 dpf demonstrated significant occlusion, with rescue of the *f5*^*-/-*^ phenotype in 73% of transgene-injected embryos (Figure 3D, p <.0001 by the Mann Whitney *U* test).

**Figure 3.**
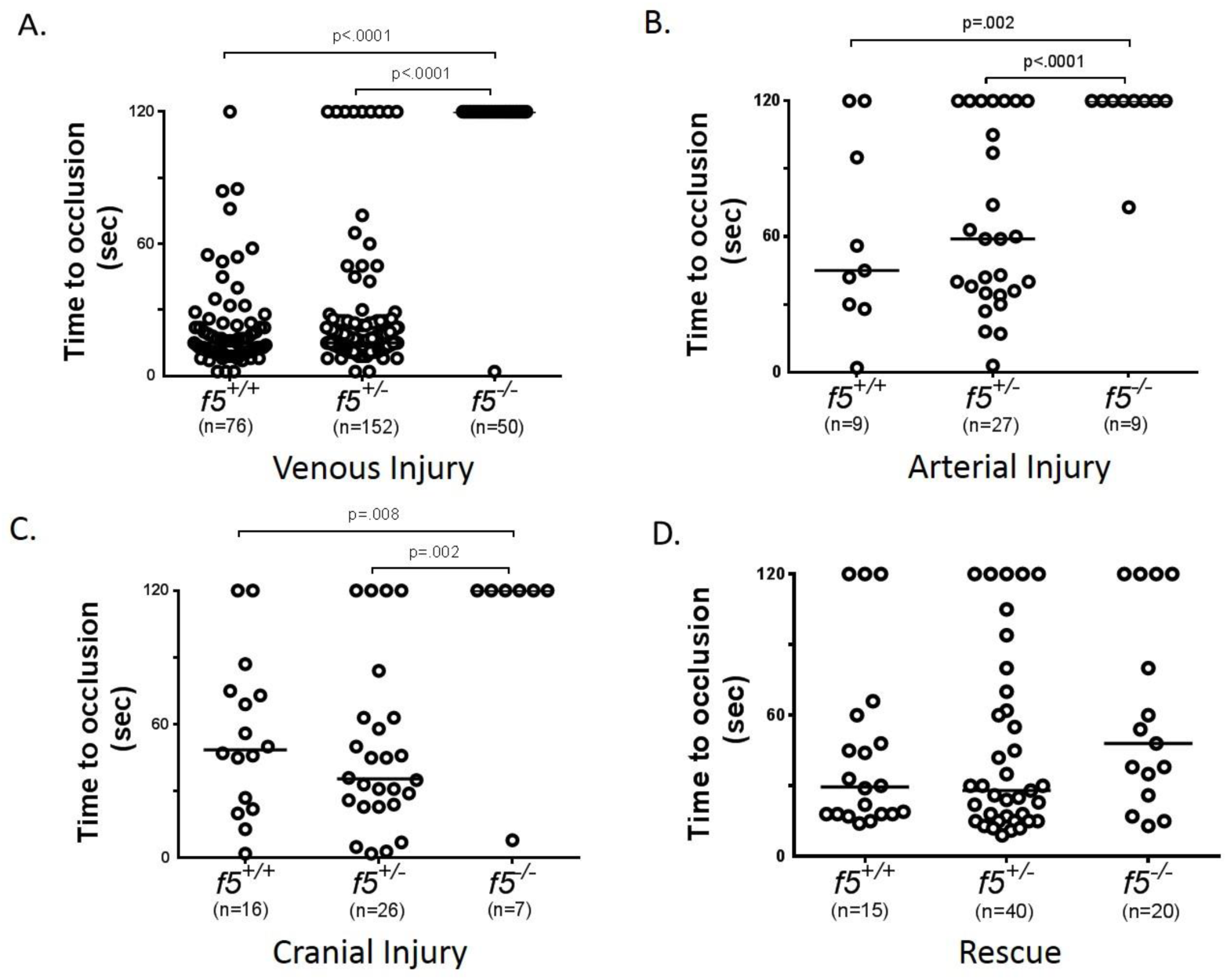
Effects of F5 deficiency on hemostasis in the venous and arterial circulation. Laser-mediated endothelial injury of the PCV was performed on larvae at 3 dpf (A). The time to occlusion was significantly prolonged in *f5*^*-/-*^ larvae in comparison with *f5*^*+/-*^ and *f5*^*+/+*^ siblings (p<0.0001, Mann-Whitney *U* test). Endothelial injury was also performed on the dorsal aorta at 9 dpf, and PMBC at 6-7 dpf with similar results (B, C, respectively). Injection of wild type zebrafish *f5* cDNA under control of the zebrafish ubiquitin promoter into 1-cell stage embryos resulted in significant rescue of the PCV hemostatic defect at 3 dpf in 73% of *f5*^*-/-*^ larvae when compared to uninjected mutants (D, p<0.001).

### Inhibition of fibrinolysis has no significant effect on the *f5*^*-/-*^ larval hemostatic defect

Given the unexpected delayed lethality of homozygous mutants as compared to a murine model, we suspected the possibility of alternate pathways for the production of fibrin. To further investigate this, we evaluated whether treatment with an antifibrinolytic, ε-aminocaproic acid, could restore venous occlusion in homozygous mutants. Previously, we demonstrated ε-aminocaproic acid mediated reversal of the hemostasis defect in fibrinogen-deficient *at3* mutants^25^, but not in *f10*^*-/-*^ larvae. In *f5* homozygous mutants, laser mediated endothelial injury revealed that treatment with ε-aminocaproic acid resulted in no statistically significant rescue (Figure 4).

**Figure 4.**
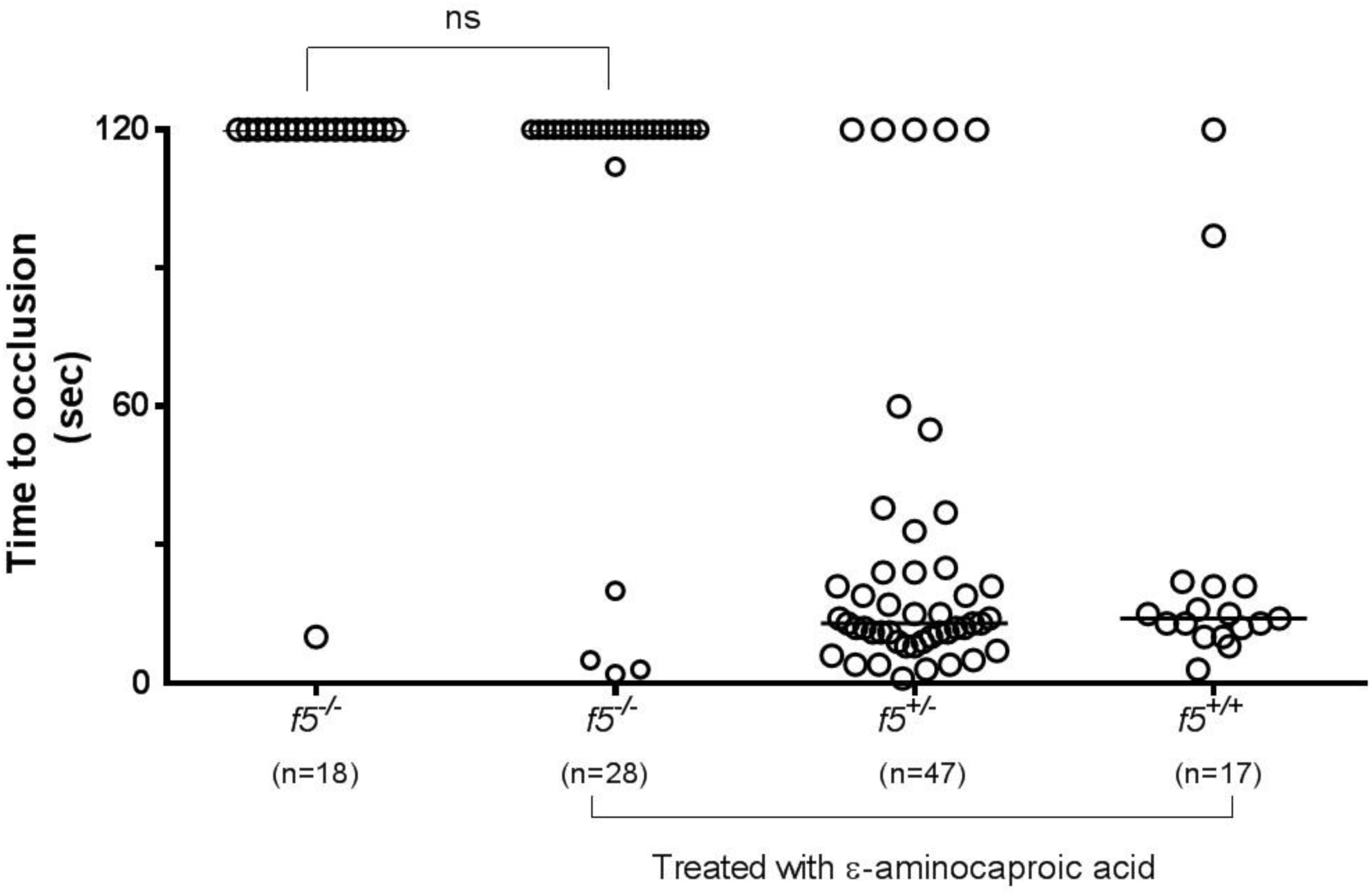
Anti-fibrinolytic treatment with ε-aminocaproic acid does not restore hemostasis in *f5* mutants. Offspring from *f5*^*+/-*^ incrosses were treated with ε-aminocaproic acid at 1 dpf and tested for venous occlusion in response to laser mediated endothelial injury at 3 dpf, followed by genotyping. There was a slight increase in the percentage of *f5*^*-/-*^ embryos that formed an occlusive thrombus (14.3% vs 5.5%) compared to untreated clutchmates, but this was not statistically significant. Horizontal bars represent the median time to occlusion. ns, not significant; sec, second.

**Figure 5.**
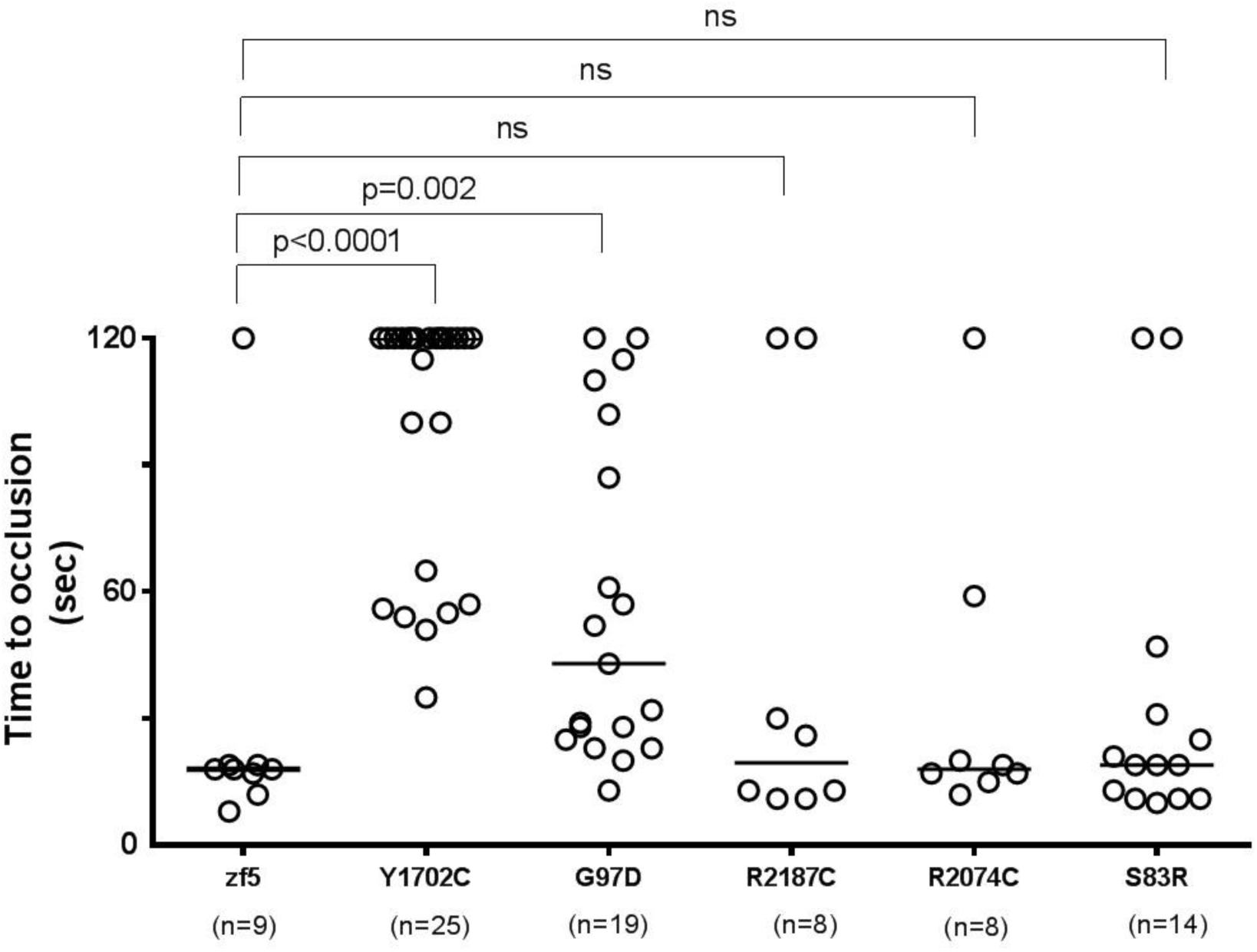
*In vivo* functional evaluation demonstrates variable severity of defects in VUS. Human F5 variants were engineered into the orthologous positions in the zebrafish *f5* cDNA under the control the of the *ubiquitin* promoter. Expression vectors were injected into 1-cell stage embryos from *f5* heterozygous incrosses. Laser mediated endothelial injury was performed at 3 dpf and time to occlusion recorded, followed by genotyping. Although Y1702C and G97D demonstrated residual occlusion, they were unable to effectively rescue (p<0.0001, p=0.002, respectively by the Mann-Whitney *U* test). Conversely, S83R, R2074C, and R2187C did not show statistically significant differences from wild-type rescue (ns, not significant by the Mann-Whitney *U* test). Horizontal bars represent the median time to occlusion. Numbering reflects the human amino acid positions. sec, second, ns, not significant.

### Evaluation of variants of unknown significance (VUS) in humans with F5 deficiency through *in vivo* evaluation in zebrafish *f5*^-/-^ mutants

With widespread availability and decreasing costs of sequencing, increasing numbers of VUS are being identified. We previously demonstrated proof of concept for the use of zebrafish for this purpose, and identified the causative mutations in patients with F10 deficiency^36^. We evaluated five *F5* mutations that had been identified in patients with clinical diagnoses of F5 deficiency (Table 2). All five of the affected amino acids are conserved across species, thus we engineered the variants into the zebrafish *f5* cDNA expression vector. These mutant plasmids were injected into zebrafish embryos from *f5* heterozygous incrosses at the 1-cell stage, followed by laser mediated endothelial injury at 3 dpf. Wild type *f5* expression plasmid, as well as the S83D, R2187C, and R2074C substitutions, were able to reverse the hemostatic defect. G97D demonstrated some activity, but was still significantly reduced from wild type, while Y1702C showed almost no activity, correlating with the phenotypes observed in patients.

**Table 2.** Human *F5* variants and clinical presentation

| Human mutation | Human amino acid variant | Orthologous zebrafish amino acid | F5 antigen | F5 activity | Allelic status | Symptoms | Reference |
| --- | --- | --- | --- | --- | --- | --- | --- |
| 5279A>G | p.Y1702C | Y1596 | 1% | <1% | Homozygous | “bleeding tendency” | (Montefusco et al., 2003) |
| 79109C>T | p.R2187C | R2092 | N/A | 10% | Compound heterozygosity with G97D | No bleeding history | unpublished |
| 32688G>A | p.G97D | G116 | N/A | 10% | Compound heterozygosity with R2187C | No bleeding history | unpublished |
| 21193C>A | p.S83R | S102 | 23% | 4% | Homozygous | N/A | unpublished |
| 6394C>T | p.R2074C | R1965 | 1-2.3% | 4% | Homozygous | Easy bruising | (Duga et al., 2003) |

## DISCUSSION

We report targeted mutagenesis of zebrafish *f5* using CRISPR/Cas9 mediated genome editing. Given early lethality due to massive hemorrhage in murine models, we hypothesized that there would be significant bleeding during the embryonic/larval stages. However, similar to F10 deficient zebrafish^36^, a large percentage of our homozygous mutants survived into adulthood. In contrast to the *f10* mutants which demonstrated a median survival of 1-2 mpf, *f5* homozygotes demonstrated progressive loss with a median survival of 4 mpf. This might be expected given the role of F5 as a cofactor in F10-mediated conversion of prothrombin to thrombin. Absence of a cofactor might be better tolerated than absence of the protease, as the former condition could retain prothrombinase activity, albeit at significantly lower efficiency. In contrast to *f10*^*+/-*^ mutants, fish heterozygous for the *f5* mutation demonstrated decreased survival compared to their wild type counterparts.

This is the third example of long term survival of a coagulation factor knockout in fish when compared to its murine counterpart. Previously, we demonstrated that genome editing of zebrafish *at3*^25^ and *f10*^36^ both result in survival into adulthood, despite mid-gestation embryonic and/or neonatal lethality in mice. Examination of the *f5*^*-/-*^ mutant larval vasculature, including arterial and venous vessels, demonstrated the inability to form thrombi in response to endothelial injury. This suggests a nearly complete hemostatic defect, so we speculate that the prolonged survival could be due to one or more of several mechanisms. Lethality in mice often occurs in the neonatal period, with bleeding occurring secondary to birth trauma. Yolk based development in the laboratory aquatic environment likely poses fewer hemostatic challenges than those encountered through the birthing process and terrestrial environs. Systemic blood pressures in mice are much higher than zebrafish^49,50^ and this may also play a role in differential risk of bleeding. Although the coagulation cascade is highly conserved between fish and mammals^31,51^, it is possible that fish have evolved unique factors or pathways that enable greater tolerance of severe bleeding defects.

We have previously observed that the hemostatic defect in At3 coagulation factor deficient larvae could be partially reversed by treatment with the fibrinolysis inhibitor ε-aminocaproic acid, but not in *f10* mutants^25 36^. These data suggest that in the former case there is enough residual fibrinogen and fibrin formation for there to be an effect, and the latter is consistent with what appears to be a complete block in fibrin production. We hypothesized that since F10 is a protease and F5 is a cofactor, we might see evidence for a low level of fibrin formation in the *f5* mutants, given that the protease is still present. This was supported by the longer term survival observed for the *f5* homozygous mutants as compared to the *f10* knockout. However, the results of fibrinolysis inhibition on induced thrombus formation were the same for both of these common pathway mutations. These data suggest that at least in the short term of the endothelial injury assay, there is not enough fibrin generated to form an occlusive thrombus.

The ability to evaluate specific human mutations for *in vivo* phenotypic rescue is an invaluable tool for determining whether novel mutations are deleterious. As the promise of whole genome and whole exome sequencing has become reality, increasing numbers of VUS have been discovered, leaving clinicians with difficulty in interpreting significance. Various databases and common resources are available to aid in this endeavor but do not provide a complete picture. *In vivo* evaluation of such mutations provides an opportunity for further characterization of the expected phenotype. Our ability to evaluate specific, clinically identified mutations is dependent on conservation across species, but this does not pose a significant hindrance as most deleterious mutations occur within well conserved regions^52^. This evaluation is particularly useful within the field of coagulation where pre-analytic variables and coincident coagulation perturbations can influence testing. Evaluation of VUS within the *F5* locus can help to identify causative mutations, as well as provide a more nuanced estimation of an individual’s patient’s bleeding phenotype. This is exceptionally advantageous in F5 deficiency, in which factor levels do not correlate well with bleeding phenotype. Further evaluation of mutations in this way may help to provide explanations for the differing phenotypes seen between patients with similar factor levels.

We evaluated five mutations, two of which had previously been described and investigated, and three VUS which were predicted to be deleterious, all of which impact conserved residues from humans to zebrafish. Y1702C is a well described missense mutation within the A3 domain which causes protein instability and may interfere with formation of disulfide bridges^53–55^. Homozygosity results in severe F5 deficiency (F5 antigen of 1% and activity of <1%^53^ and no significant rescue was observed in our evaluation. G97D (A1 domain) and R2187C (C2 domain) have not been previously described but were found to be compound heterozygous in a patient with an F5 antigen level of 10%. G97D was not able to restore venous occlusion in our analysis, although R2187C was able to do so, implying that the latter is a less deleterious mutation which may compensate clinically. S83R (A1 domain) has not previously been described and was found to be homozygous in a patient with F5 antigen and activity levels of 23% and 4%, respectively. Despite its low activity, it was able to effectively restore venous occlusion. R2074C (also in the C2 domain) results in decreased secretion of F5 and enhanced intracellular degradation, and homozygosity results in low but detectable F5 activity (4%)^56^. Venous occlusion was restored despite this defect, demonstrating that this molecule does retain *in vivo* coagulant activity and strategies that mitigate the secretion defect could be exploited as a clinical strategy to restore hemostasis in affected patients.

All of the mutations demonstrated some ability to enhance occlusion, even when statistically different from wild type F5, demonstrating that as in mammals, low levels of F5 activity may be sufficient for augmentation of thrombin generation in zebrafish. This is consistent with the relatively mild clinical bleeding phenotypes observed, even in patients with unmeasurable F5 activity. Half normal endogenous thrombin potential (ETP) has been shown with as little as 1% F5, with absent or mild bleeding found in patients with ETPs of 30% of higher^17^. Interestingly, our rescue results do correspond with both bleeding symptomatology and reported factor activity levels. Historically, activity level has had limited correlation with severity of bleeding^57^. Our *in vivo* evaluation could be increasingly beneficial in patients for whom factor activity levels do not correlate with bleeding phenotype, ideally performed early on and providing treatment and prognostic guidance. As patients with identical activity levels may have quite different bleeding phenotypes, this tool may prove advantageous in providing a more nuanced view of individual hemostasis. The fecundity of zebrafish and ease of working with large numbers allow for easily repeated experiments which may ultimately unveil hemostatic differences which would otherwise remain undiscovered.

In summary, our model indicates strong conservation between zebrafish and mammalian F5, including the site of its synthesis and necessity for hemostasis. Surprisingly, embryos and larvae tolerate what is a severe and lethal defect in mammals, allowing further studies not available in the mouse. This viability, now across 3 coagulation factor mutants with important roles in the common pathway, suggests the possibility of species-specific factors enabling survival. Further studies to identify such factors or other genetic modifiers could lead to novel diagnostic and therapeutic modalities for patients with bleeding or clotting disorders.

## Acknowledgements

This study was supported (in part) by research funding from an NHF/Shire Clinical Fellowship award, an HTRS/Novo Nordisk Mentored Research Award in Hemophilia and Rare Bleeding Disorders from the Hemostasis and Thrombosis Research Society (HTRS), which was supported by an educational grant from Novo Nordisk Inc. and NIH T32 HL007622, and NIH K12 HD028820 (A.W.); NIH T32 HL125242 (S.J.G.); National Hemophilia Foundation Judith Graham Pool Award (M.S.R.); Ricerca Corrente of the Italian Ministry of Health (F.P.); NIH R01 HL124232 and HL125774 and Bayer Hemophilia Awards Program (J.A.S.). J.A.S. is a recipient of the American Society of Hematology Scholar Award and is the Diane and Larry Johnson Family Scholar of Pediatrics and Communicable Diseases.

## Authorship Contributions

Contribution: A.C.W. designed and performed research, analyzed data, and wrote the manuscript; S.G., K.L. and M.R. designed and performed research and analyzed data; A.C.F and C.R. performed research and analyzed data; R.A., S.D., M.M. and F.P. designed research; R.A. and S.D. genetically characterized F5-deficient patients; J.A.S. designed and supervised research, analyzed data, and edited the manuscript.

